# The ATP synthase subunit β (ATP5B) is an entry factor for the hepatitis E virus

**DOI:** 10.1101/060434

**Authors:** Zulfazal Ahmed, Prasida Holla, Imran Ahmad, Shahid Jameel

**Affiliations:** Virology Group, International Centre for Genetic Engineering and Biotechnology, New Delhi, India

## Abstract

Hepatitis E occurs sporadically and as outbreaks due to contamination of drinking water. The causative agent, hepatitis E virus (HEV) is a hepatotropic non-enveloped RNA virus, which grows poorly *in vitro.* Consequently, many aspects of HEV biology are poorly characterized, including its cellular receptor and entry mechanism(s). Previous studies from our laboratory have shown that heparan sulfate proteoglycans (HSPGs) act as attachment factors for the virus. In the absence of purified high titer infectious virus, we have used hepatitis E virus-like particles (HEV-LPs) expressed and purified from *E. coli* to identify HEV entry factor(s) on liver cells in culture. Using a pull down and mass spectrometric approach, we identified the ATP synthase subunit β (ATP5B) to bind the HEV capsid protein. Its role in the entry of HEV was then validated using antibody and siRNA mediated approaches, and infectious HEV from the stools of a hepatitis E patient. Though ATP synthase is largely a mitochondrial protein, the cell surface expressed form of ATP5B is implicated in other viral infections.

## INTRODUCTION

Hepatitis E virus (HEV) causes an acute and generally self-limited viral hepatitis with a mortality rate of 0.5-4% in the general population, which can be as high as 20-30% in infected pregnant women (Khuroo, 2011). The virus is enterically transmitted and several large outbreaks have occurred in Asia and Africa due to contamination of drinking water with fecal matter. In developed countries, HEV is transmitted primarily through the consumption of raw or undercooked meat from infected animals, with pigs being an important reservoir (Meng, 2011; Sonoda et al., 2004; Yugo & Meng, 2013).

The HEV is classified as a Hepevirus in the family *Hepeviridae* (Smith et al., 2014). It is a 27-34 nm non-enveloped virus with a ~7.2 kb positive sense polyadenylated and capped RNA genome (Emerson & Purcell, 2006). Short untranslated regions (UTRs) flank the coding region of the genome, which includes three open reading frames (ORFs) called *orf1, orf2* and *orf3* (Emerson & Purcell, 2006). Of these, *orf2* encodes the viral capsid protein (Jameel et al., 1996). Efficient cell culture systems and suitable small animal models are not available for the propagation of HEV, due to which details of HEV biology are limited to the role of individual ORFs and the proteins encoded by these through subgenomic or replicon expression strategies. The receptor(s) for HEV entry have also not been identified thus far. *In vitro* transcribed and capped viral genomic RNA is infectious for some cultured cells and non-human primates following transfection and intrahepatic injection, respectively. However, the infectivity and cell-to cell-spread of the virions thus generated is poor (Tanaka et al., 2007).

The structure of HEV has been studied by expressing the ORF2 capsid protein in various surrogate systems, such as insect cells, mammalian cells and *E. coli* (Zheng et al., 2010; Zhang et al., 2001). Of these, p239 (aa 368-606), which contains the surface localized protruding domain, when expressed in *E. coli*, folds into 23 nm virus-like particles with a T=1 symmetry that antigenically and structurally resemble HEV (Ahmad et al., 2011). Further, in human clinical trials p239 has shown efficacy as a preventive vaccine (HEV 239) against hepatitis E (Li et al., 2005; Zhu et al., 2010).

The initial attachment of many viruses to host cells takes place via cell surface glycolipids or glycoproteins (de Haan et al., 2005; Vlasak et al., 2005). This is a relatively non-specific electrostatic interaction that allows viral particles to be concentrated on the cell surface. Following initial attachment, most viruses bind to high affinity receptor(s) leading to entry and productive infection (Grove & Marsh, 2011). The nature of the molecules(s) utilized by a virus to bind and enter cells determines its tropism, internalization, uncoating and trafficking to sites of intracellular replication. Some viruses bind single receptors, such as CD155 for the poliovirus (Mendelsohn et al., 1989) or low density lipoprotein receptor (LDLR) for human rhinovirus 2 (Hofer et al., 1994). Other viruses bind multiple molecules on the host cell, which usually occurs in temporally organized steps. For example, the SARS coronavirus can bind either angiotensin converting enzyme (ACE) or liver-SIGN (L-SIGN) to gain entry into cells (Li et al., 2003; Jeffers et al., 2004). The human immunodeficiency virus (HIV) binds two distinct molecules on the cell surface, CD4 as the primary receptor (Dalgleish et al., 1984; Klatzmann et al., 1984) followed by either CCR5 or CXCR4 as the co-receptor (Choe et al., 1996; Deng et al., 1996), both interactions being essential for productive infection of target cells by HIV. The hepatitis C virus (HCV) uses multiple entry factors-Scavenger receptor-B1 (SR-B1) (Scarselli et al., 2002), CD81 (Pileri et al., 1998), tight junction proteins Claudin-1 (CLDN1) (Evans et al., 2007), Occludin (OCLN) (Ploss et al., 2009) and other molecules like EGFR and EphA2 (Lupberger et al., 2011), Transferrin receptor (Martin and Uprichard, 2013) and Niemann-Pick C1-like 1 cholesterol absorption receptor (NPC1L1) (Sainz et al., 2012). Enveloped viruses bind target cells through their surface proteins embedded in the lipid envelopes, while non-enveloped viruses bind via their capsid protein(s). In either case, the binding of viruses to receptors triggers signalling leading to activation of endocytosis or modulation of actin dynamics for lateral movement and clustering of the virus on the plasma membrane prior to entry (Lehmann et al., 2005; Burckhardt & Greber, 2009). It can also directly trigger conformational changes in the virus structure to an unstable intermediate that releases the genome. Following receptor engagement, some enveloped viruses (e.g. HIV) undergo fusion and non-enveloped viruses (e.g. poliovirus) undergo uncoating at the cell surface (Kelian & Rey, 2013; Sieczkarski & Whittaker, 2005; De Sena & Mandel, 1977). Most viruses, however, are internalized by endocytosis into endosomal compartments and reach their site of replication by escaping from endosomes. The trigger for this process can be low pH or proteolysis of the capsid by proteases in early or maturing endosomes (Marsh & Helenius, 2006). Examples of this include acidification of the influenza virus core due to H+ translocation through its M2 protein (Wharton et al., 1994), and cleavage of the Ebola virus glycoprotein by Cathepsin to reveal determinants that bind the endosomal receptor NPC-1, allowing translocation to the cytosol (Chandran et al., 2005; Cote et al., 2011; Carette et al., 2011).

Previous studies from our laboratory have shown that HEV binds liver cells via heparin sulphate proteoglycans (HSPGs) present on cell surface Syndecans (Kalia et al., 2009). We have recently used the HEV p239 VLP expressed in *E. coli* and virus infection to show that HEV uses a Clathrin, Dynamin-2 and membrane cholesterol dependant entry pathway in liver cells (Holla et al., 2015). Here, we use the same VLPs to pull down proteins in purified membrane fractions from permissive (Huh-7) and non-permissive (HEK293) cell lines. This approach and further validation using antibody and siRNA-mediated inhibition, fluorescent imaging and virus infection identified that ATP synthase subunit β (ATP5B) is involved in the entry of HEV. This is the first study to identify an entry factor for HEV.

## MATERIALS AND METHODS

**Cells and reagents.** Huh-7, PLC/PRF/5, HeLa, HEK293 and A549 cells were grown in DMEM supplemented with 10% Fetal Calf Serum (FCS), 100 U/mL Penicillin and 100 μg/mL Streptomycin (all from Gibco, Life Technologies). PCR primers were designed manually and obtained from Sigma Aldrich (Bengaluru, India). Phusion Hi-Fidelity DNA polymerase was from Finnzymes (Thermo Fischer Scientific, Inc.), restriction enzymes and calf intestinal phospahtase (CIP) were from New England Biolabs Inc., and DNA Ligase was from Promega Corporation. The Nickel-NTA Superflow beads were obtained from Qiagen, Germany. The cell surface protein isolation kit and Zeba Desalting columns were purchased from Thermo Scientific, USA, the protease inhibitor cocktail was from Roche, and mass spectrometry grade Trypsin Gold was from Promega Corporation. On Target PLUS Smart Pool siRNAs against ATP5B were purchased from Dharmacon Inc. CTxB-biotin was from Sigma Aldrich (Bengaluru, India). Antibodies were obtained from the following sources: HEV capsid protein (Anti-HEV capsid antibody, clone 1E6) from LS Bio (USA); ATP5B (N-terminal region) from Sigma-Aldrich; Cytochrome C, Calnexin, Actin, GAPDH, MAPK/ERK and EGFR from Santa Cruz Biotechnologies Inc. (Santa Cruz, CA, USA); Alexa Fluor 488 and Alexa Fluor 568 coupled anti-rabbit and anti-mouse antibodies, and HRP-conjugated secondary anti-goat, anti-rabbit and anti-mouse IgG from Calbiochem and Millipore (MA, USA).

**Transfections.** Plasmid transfections were done with jetPRIME (Polypus Transfection) according to manufacturer’s instructions. For siRNA transfections, 0.25 ×10^6^ cells (Huh-7 or HEK293) were seeded in each well of a 6-well plate and grown without antibiotics. After reaching 70% confluence, cells were transfected with 50 nM siRNA against ATP5B or with a non-specific scrambled siRNA as control, incubated for 4 hours in a serum-free medium followed by complete medium without antibiotics for 36 hours. At this time, cells were transfected again with siRNA (to ensure maximum knockdown) and harvested after 72 hours of the first transfection. The cells were then used for either western blotting or flow cytomtery.

**Cloning, expression, purification, characterization and labelling of HEV-LP.** Cloning of the HEV-LP was done as described previously (Holla et al., 2015). Briefly, the *orf2* region corresponding to amino acids 368-606 (nucleotides 1104-1821) were PCR amplified from the pSK-HEV2 plasmid (Sar-55 strain, genotype 1) with primers containing NcoI and XhoI sites, and cloned into the pET28b vector. The positive clones were transformed into the *E. coli* expression strain BL21(DE3) and the VLP monomer was expressed as a C-terminally 6× Histidine-tagged protein. For expression, 250 ml bacterial culture was induced with 1 mM isopropyl-D-thiogalactosidase (Sigma-Aldrich) for 4 hours at 37°C. The cells were harvested by centrifugation and broken by brief sonication in lysis buffer (50 mM Tris-Cl, pH 8.0, 5 mM EDTA, 10mM NaCl, 0.5% Triton X-100, 0.1 mM phenylmethylsulfonyl fluoride and 0.1% NaNs). The cell lysate was then treated with 10 mM MgSO_4_ for 10 min at room temperature to chelate EDTA followed by 0.1 mg/ml lysozyme for 20 min at 37°C. Inclusion bodies were pelleted by centrifugation at 8,000 rpm (Sorvall SLA 300 rotor) for 15 min at 4°C, resuspended in 100 mM Tris-Cl, pH 8.0 with 100 mM glycine and sonicated for 2 min. The resuspended inclusion bodies were added dropwise to the dispersion medium (100 mM Tris pH 8.0, 50 mM glycine, 8 M urea, 5 mM reduced glutathione and 0.5 mM oxidized glutathione, and left overnight on a rocker. This preparation was then bound to Ni-NTA bead (Qiagen, Germany) for 1 hour at room temperature on a rotaspin machine. The beads were washed with binding buffer containing 20 mM imidazole and bound proteins were eluted in binding buffer containing 200 mM imidazole. The eluted protein was concentrated using a 10 kDa Amicon filter (Millipore), subjected to refolding by rapidly diluting 40-fold in a refolding buffer containing 1 mM EDTA, 5 mM reduced glutathione, 0.5 mM oxidized glutathione, 0.5 mM Arginine and 20 mM ethanolamine, pH 8.0, for 36 hr at 4°C. The refolded protein was concentrated again using a 10 kDa Amicon filter and dialyzed against PBS. Electron microscopy and FITC labeling of the HEV-LP have been described elsewhere (Holla et al., 2015).

**Surface protein isolation.** Cell surface proteins were isolated with two different methods. The first used the Total Cell Surface protein isolation Kit (Thermo Scientific) with some modifications. Briefly, cells were grown in 100 mm dishes up to 80% confluence, washed and treated with Sulfo-NHS-SS-Biotin in PBS for 30 min on ice. After washing, the cells were treated with Quenching Solution, pelleted and lysed in PBS containing the protease inhibitor cocktail by sonication at 25 amp pulse for 5 sec (and a 10 sec off interval) for a total of 3 min on ice. The lysate was incubated with NeutraAvidin agarose beads for 2 hr at 4°C with end-to-end mixing. The beads were washed with PBS and bound biotinylated proteins were eluted in PBS containing 50 mM DTT, which was subsequently removed by dialysis against PBS. In the second approach, membrane fractions were isolated by ultracentrifugation. Cells in 100 mm dishes were washed with PBS, scraped and lysed in PBS containing protease inhibitor cocktail as above. The cell debris and nuclei were removed by centrifugation at 600 × g for 10 min at 4°C, and the membrane fraction was pelleted down in a SW41Ti rotor (Beckman) at 29,000 rpm for 1 hr at 4°C. This membrane pellet was washed once with PBS containing protease inhibitor cocktail by ultracentrifugation as above and resuspended in 400 μl PBS. This was layered over a sucrose step gradient made up of 900 μ each of 45%, 35%, 20% and 5% sucrose (w/v) up to a total volume of 4 ml. The gradient was centrifuged at 42,000 rpm in a SW55-Ti rotor for 16 hr at 4°C. From this, nine fractions of 330 μl were collected from the top (F1-F9) and 40 μl of each fraction was resolved on a SDS-12% polyacrylamide gel, transferred to nitrocellulose membrane and western blotting was done for ATP5B, Calenexin, Cytochrome c and EGFR.

**Mass spectrometric identification of membrane proteins.** To identify HEV-LP interacting membrane proteins, the fractions from Huh-7 and HEK293 cell lines were pre-cleared with Ni-NTA beads for 2 hr before incubating with purified HEV-LP overnight at 4°C on a rotator. The mixture was then added to Ni-NTA beads and incubated at room temperature for 1 hr. The beads were washed and proteins were eluted by boiling in 1× SDS loading dye at 95°C for 10 min. Proteins were separated by SDS-PAGE and stained with Coomassie Blue. The bands were excised, cut into small pieces and in-gel tryptic digestion was carried out as described (Shevchenko et al., 2006). Briefly, gel pieces were destained in 100 μl of destaining solution (25 mM NH4HCO3 and 50% CH3CN) by rinsing 3-4 times, centrifuging at 1000 rpm for 1 min and discarding the supernatant till the gel pieces were completely destained. These were then dried in a vacuum concentrator. Trypsin solution (13 ng/μl, in 25 mM NH_4_HCO_3_) was added to the dried gel pieces and incubated at 37°C overnight, following which the sample was centrifuged at 1000 rpm for 1 min and digested peptides were extracted in 20 μ of extraction solution (75% CH3CN and 5% Trifluoroacetic acid) by vortexing for 30 min. The supernatant was collected, the extraction procedure was repeated 3-4 times, and the pooled extracts dried in a vacuum concentrator before submitting for mass spectrometric analysis. For the insol tryptic digestion, total cell surface fractions from Huh-7 and HEK293 cells were isolated according to the kit protocol. The final eluate, which contained 50 mM DTT, was mixed with 6 volumes of UA buffer (8 M urea in 0.1 M Tris/HCl pH 8.5) and added onto a 30 kDa cut-off Nanosep centrifugal filter (Pall Corporation) and centrifuged at 14,000 × g for 15 min. Three volumes of IAA solution (0.05 M iodoacetamide in UA buffer) were added to the filter for 20 min, followed by four washes with UA buffer. The filter was then equilibrated for tryptic digestion by incubating with ABC solution (0.05 M NH4HCO3) for 20 min, and centrifuging at 14,000 × g. This step was repeated twice before adding trypsin diluted in ABC buffer (trypsin to protein ratio 1:100) for 8 hr at 37°C. The digested peptides were eluted in a new collection tube, acidified with trifluoroacetic acid (TFA) and subjected to mass spectrometry.

**Immunofluorescent staining and confocal microscopy.** For all microscopy experiments, cells were grown on coverslips in 12-well plates. To stain intracellular molecules, cells were fixed with 2% paraformaldehyde in PBS for 15 min and then permeabilized with 0.04% Triton X100. Blocking was done with 5% BSA for 30 min at room temperature followed by staining with appropriate primary antibodies and fluorophore-conjugated secondary antibodies in 1% BSA for 1 hr. After each antibody incubation, cells were washed five times with PBS. Coverslips were mounted onto slides with ProLong Gold Antifade Reagent with DAPI. To study the colocalization of internalized FITC-VLP and ATP5B, the FITC-VLPs were bound to cells on ice for 1 hr, washed with ice-cold PBS to remove unbound material and then shifted to 37°C to allow internalization for different times. Internalization was terminated by adding ice-cold PBS to cells. Surface bound VLPs were removed by first treating cells with trypsin-EDTA for 30 sec and then with surface stripping buffer (0.2% glacial acetic acid, 500 mM NaCl) for 1 min. The cells were fixed, permeabilized and stained with ATP5B primary antibody and Alexa Fluor 568 conjugated secondary antibody. All images were acquired on a Nikon A1R confocal laser scanning microscope (cLSM). Images were processed using Image J (http://rsbweb.nih.gov/ij/download.html) and colocalization coefficients were calculated using the NIS-Elements Microscope Imaging Software.

**Plasmid DNA isolation.** All plasmids were prepared using the RBC HiYield Plasmid DNA kit (Real Biotech Corporation, Taiwan) or cesium chloride density gradient centrifugation using standard protocols (Sambrook et al., 1989).

**Flow cytometry.** To evaluate the binding of HEV-LPs or cell surface levels of ATP5B by flow cytometry, cells were scraped, washed twice with FACS buffer (PBS + 0.1% FCS), blocked and incubated on ice for 1 hr. Cells were then incubated with either HEV-LP for 1 hr on ice or with anti-ATP5B polyclonal antibody in FACS buffer for 2-3 hr at 4°C, and washed three times with FACS buffer. The HEV-LP bound cells were stained with anti-HEV capsid followed by Alexa Fluor 488 conjugated secondary antibody; the anti-ATP5B stained cells were further stained with an appropriate secondary antibody. Cells were washed and subjected to flow cytometric analysis. Appropriate isotype control antibodies were used as primary antibodies for all stainings.

For antibody and HEV-LP binding and blocking experiments, cells were incubated with anti-ATP5B (or isotype control) antibodies for 2 hr on ice or with HEV-LPs for 1 hr on ice. After incubation, the cells were fixed, washed with FACS buffer and incubated with HEV-LPs for 1 hr or with ATP5B antibodies for 2 hr, respectively. Cells were then washed and stained with anti-HEV capsid antibody and a fluorophore conjugated secondary antibody and subjected to flow cytometry.

All flow cytometric measurements were carried out on a CyAn ADP flow cytometer (Dako Cytomation). Data were analysed using the Summit Software (Dako Cytomation) and histograms were made using FlowJo (Tree Star, Inc.). Data were plotted as a percentage of fluorescent cells relative to mock (DMSO or other) treated cells.

**Western blotting.** Isolated membrane proteins were run on 10 or 12% denaturing polyacrylamide gels containing SDS, transferred to nitrocellulose membranes (Advanced Microdevices, India), blocked with 5% w/v BSA for 2 hr at room temperature and probed with various primary (overnight at 4°C) and HRP conjugated secondary antibodies (30 min at room temperature) diluted in TBS-T containing 1% BSA. Blots were visualized using Immuno-Cruz Western Blotting Luminol Reagent (Santa Cruz Biotechnologies Inc.) and exposed to X-ray films.

**HEV infection and estimation of viral titres.** The stool of a HEV-infected patient was used to make a 10% suspension in PBS, which was clarified by centrifugation at 10,000 × g for 20 min at 4°C. This was then filtered successively through 0.45 μm and 0.22 μm filters and stored in single-use aliquots at −80°C. To estimate titres, RNA was isolated from 150 μl of this suspension using the QIAamp Viral RNA kit (Qiagen) and subjected to one step RT-Real time PCR (SuperScript III Platinum One-Step qRT-PCR Kit w/ROX, Life Technologies) using HEV specific primers and probes (Jothikumar et al., 2006). A standard curve was generated by this method using known concentrations of *in vitro* transcribed HEV RNA. Cells were pre-treated with anti-ATP5B (or isotype control) antibodies for 1 hr, followed by infection with 2 million genome equivalents of the stool suspension per million cells for 5 hr, followed by three PBS washes. The cells were then cultured in virus maintenance medium (DMEM containing 2% FCS and 2 mM MgCl_2_) for 40 hr. Total cellular RNA was isolated using TriZol (Invitrogen) and subjected to qRT-PCR (described above) to determine titres.

**Lipid raft isolation.** Lipid rafts were isolated according to published protocols (Legler et al., 2005). Briefly, FITC-VLP and CtxB-Biotin were bound on ice to Huh-7 cells for 1 hr and lysed in chilled MNE buffer (25 mM MES, pH 6.5, 150 mM NaCl, 2 mM EDTA) containing 1% Triton X100 using a Dounce homogenizer. Alternatively, cells were washed and shifted to 37°C for 1 hr after binding of FITC-VLP and CTxB-biotin. The lysate was mixed with an equal volume of 90% sucrose in MNE buffer and placed in the bottom of an ultracentrifuge tube. This was overlaid with two volumes of 35% sucrose followed by 5% sucrose. The gradient was centrifuged at 175,000 × g for 20 hr in a SW41Ti rotor (Beckman). One mL fractions were collected from the top and subjected to western blotting for FITC-VLP and EGFR, the latter as a marker of non-raft fractions. To probe for raft-enriched fractions, these were blotted on nitrocellulose membrane, blocked in 5% BSA/PBS/0.1% Tween 20 and probed with Streptavidin-HRP at a dilution of 1:5000.

## RESULTS

**Preparation of HEV-LPs and identification of binding partners.** The region corresponding to amino acids 368-606 of the HEV capsid protein (p239) was cloned and expressed in *E. coli* as described in Materials and Methods. Increasing amounts of the purified protein were run on a reducing SDS-polyacrylamide gel, which migrated between the 26 and 34 kDa markers (Fig. 1a). The purified protein also assembled into 20-40 nm virus like particles, as visualized by electron microscopy (Fig. 1b); these are called the hepatitis E virus-like particles (HEV-LPs). Dynamic light scattering (DLS) studies and gel permeation chromatography were also used to characterize the HEV-LPs as detailed elsewhere (Holla et al., 2015). The purified protein eluted in the void volume of a TSK3000 silica gel column (separation range 10-500 kDa), indicating that higher order structures were being formed, and DLS measurements showed the average diameter of the particles to be 34 nm. We then checked the ability of these HEV-LPs to bind to cells of hepatic (Huh-7) and non-hepatic (HEK293) origin by flow cytometry, and found the binding to be higher on the former compared to the latter (Fig. 1c). These cell lines are considered permissive and non-permissive for HEV, respectively. Additionally, compared to HEK293 cells, the binding of HEV-LPs was higher on two other cell lines that are permissive for HEV infection (Takahashi et al., 2012)-PLC/PRF/5 (hepatoma) and A549 (lung carcinoma) (Fig. S1).

**Figure 1.**
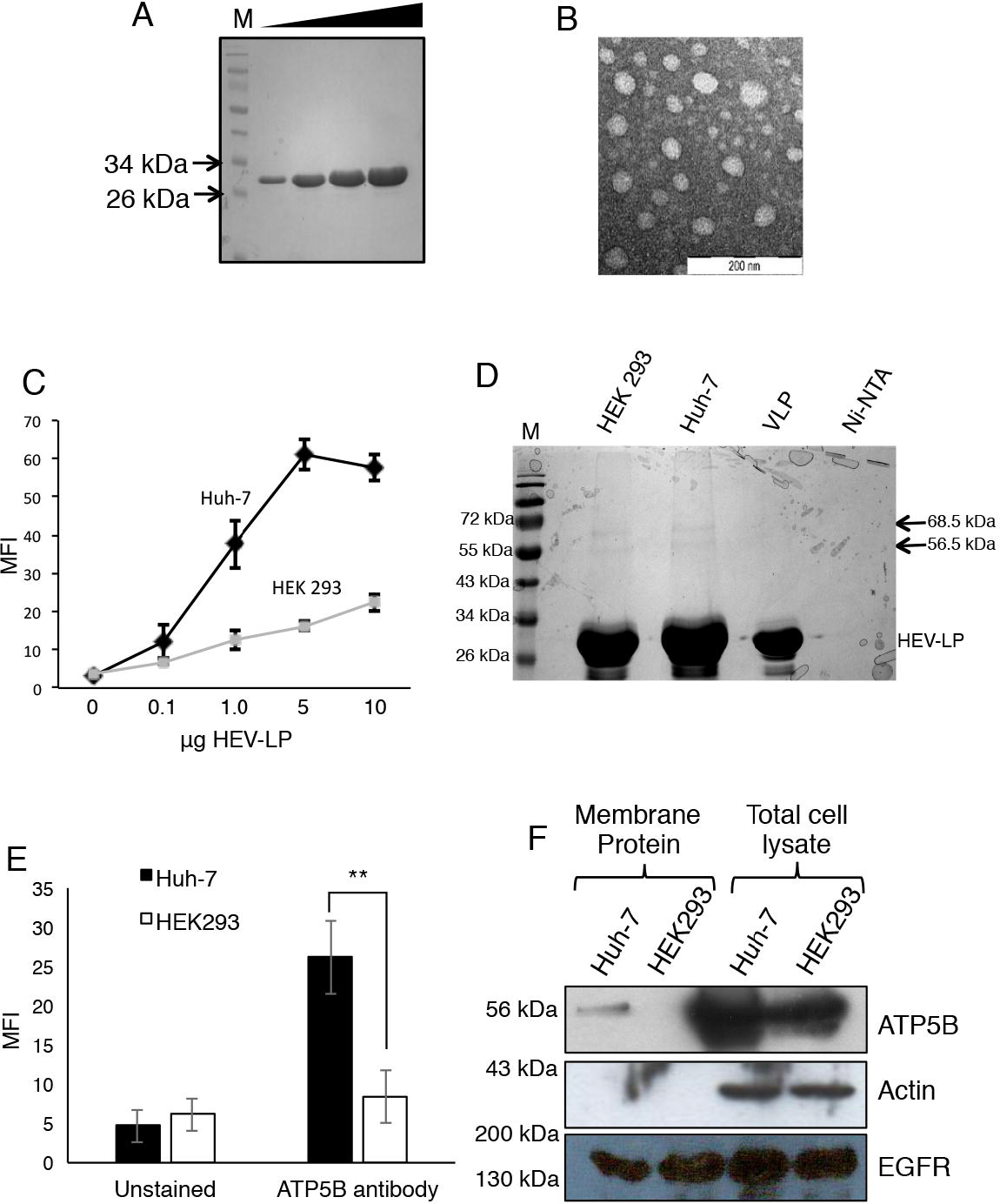
**(A)** The HEV-LP was expressed in E.*coli* BL21 (DE3) and purified from inclusion bodies. Increasing concentrations (7.5 to 30 μg) of purified recombinant protein were run on a SDS-12% polyacrylamide gel and visualized by Coomassie Blue staining. **(B)** Electron microscopic image of the HEV-LP preparation was obtained by loading purified protein onto Formvar-coated copper grids and visualizing at 97,0000× magnification. **(C)** Different amounts of HEV-LP (100 ng to10 μg) were bound to Huh-7 or HEK293 cells, stained with anti-HEV capsid antibody, and subjected to flow cytometry. Data represents mean fluorescence intensity from three independent experiments. **(D)** Membrane proteins from Huh-7 and HEK293 cells were used in a pull-down experiment to identify HEV entry factors. Eluates were separated on an SDS-polyacrylamide gel and stained with Coomassie Blue. Two bands of the indicated molecular sizes were observed. Positions of protein marker (M) bands are shown on the left. **(E)** Surface levels of ATP5B were measured on Huh-7 and HEK293 cells by staining cells with anti-ATP5B (or isotype control) antibodies followed by flow cytometry; data is representative of three independent experiments ± SD (**p<0.01). **(F)** Membrane protein fractions and total cell lysate from Huh-7 and HEK293 cells were separated on a SDS-12% polyacrylamide gel, transferred to nitrocellulose membrane and probed for ATP5B using specific antibodies. Membrane fractions were probed for Actin to check for cytoplasmic contamination and EGFR was used as a loading control. Protein markers are indicated.

The C-terminal 6X Histidine tag present on the p239 monomer was used to immobilize the HEV-LPs to Ni-NTA beads. Membrane fractions from HEK293 and Huh-7 cells were passed over the immobilized HEV-LPs, the interacting proteins were eluted and resolved by SDS-PAGE. Two bands corresponding to 56.5 kDa and 68.5 kDa were observed for both the cell lines (Fig. 1d). The bands were excised, subjected to in-gel tryptic digestion, and the proteins were identified by mass spectrometry to be ATP synthase subunit β (ATP5B) and Ribophorin I (RPN I) (Table S1).

**ATP5B is present on the surface of Huh-7 and HEK293 cells.** To ensure that ectopic (and not mitochondrial) ATP synthase was obtained from the pull-down assay, we isolated the total cell surface proteins from Huh-7 and HEK293 cells, and subjected these fractions to in-sol tryptic digestion and MALDI-TOF spectrometry. ATP5B was identified from Huh-7 cells, and not from HEK 293 cells (Table S2). Additionally, the alpha subunit (ATP5A) was also identified from Huh-7 cells, indicating that the two subunits may be forming a larger complex on the cell surface We also studied the localization and expression levels of ATP5B by flow cytometry and western blotting. Unpermeabilized Huh-7 and HEK293 cells were surface stained with ATP5B antibodies and subjected to flow cytometry. The ATP5B levels were higher on Huh-7 cells as7 compared to HEK293 cells (Fig. 1e; p<0.01), and this was consistent with HEV-LP binding to these cells (Fig. 1c). We also subjected membrane fractions from Huh-7 and HEK293 cells to western blotting for ATP5B. A band corresponding to ATP5B was observed in membrane fractions from Huh-7 and HEK293 cells, but the levels were significantly higher in the former as compared to the latter (Fig. 1f). Total cell lysates from Huh-7 and HEK293 cells showed high levels of ATP5B in both cell types. To ensure that membrane fractions were devoid of cytoplasmic contaminants, they were probed for Actin and EGFR was used as a loading control. Plasma membrane fractionation protocols are subject to contamination from internal membranes. To ensure that ATP5B was not a contaminant from internal membranes, we carried out sucrose density gradient centrifugation and probed the collected fractions with antibodies against various subcellular markers: EGFR (plasma membrane), Calnexin (endoplasmic reticulum) and Cytochrome C (mitochondria) (Fig. S2). The ATP5B protein was found quantitatively in the EGFR marked plasma membrane fractions, with no visible contamination from internal membranes (ER, mitochondria) as seen from the lack of Calnexin and Cytochrome C in these fractions from both Huh-7 and HEK293 cells.

**Blocking of ATP5B reduces HEV-LP binding and virus infection.** We then checked whether antibody-mediated masking of cell surface exposed ATP5B affected the binding of HEV-LPs. For this, Huh-7 cells were pre-incubated with anti-ATP5B antibodies on ice, followed by the binding of HEV-LP, staining with anti-capsid antibodies and quantitation of binding with flow cytometry. The pre-incubation of cells with anti-ATP5B antibodies reduced the binding of HEV-LPs compared to cells treated with an isotype control antibody (Fig. 2a, 2b; p<0.005). Conversely, when Huh-7 cells were first incubated with HEV-LPs, fixed and subsequently stained with anti-ATP5B antibodies, the detectable surface levels of ATP5B were significantly reduced (Fig. 2c, d; p<0.05). To further validate these findings, we infected antibody pre-treated Huh-7 cells with HEV and estimated the virus in these cells two days post-infection by quantitative RT-PCR as described in Materials and Methods. The viral titers were found to be reduced by 1.5-2 log10 levels in cells pre-treated with ATP5B antibodies compared to the isotype control antibody (Fig. 2e, p<0.01).

**Figure 2.**
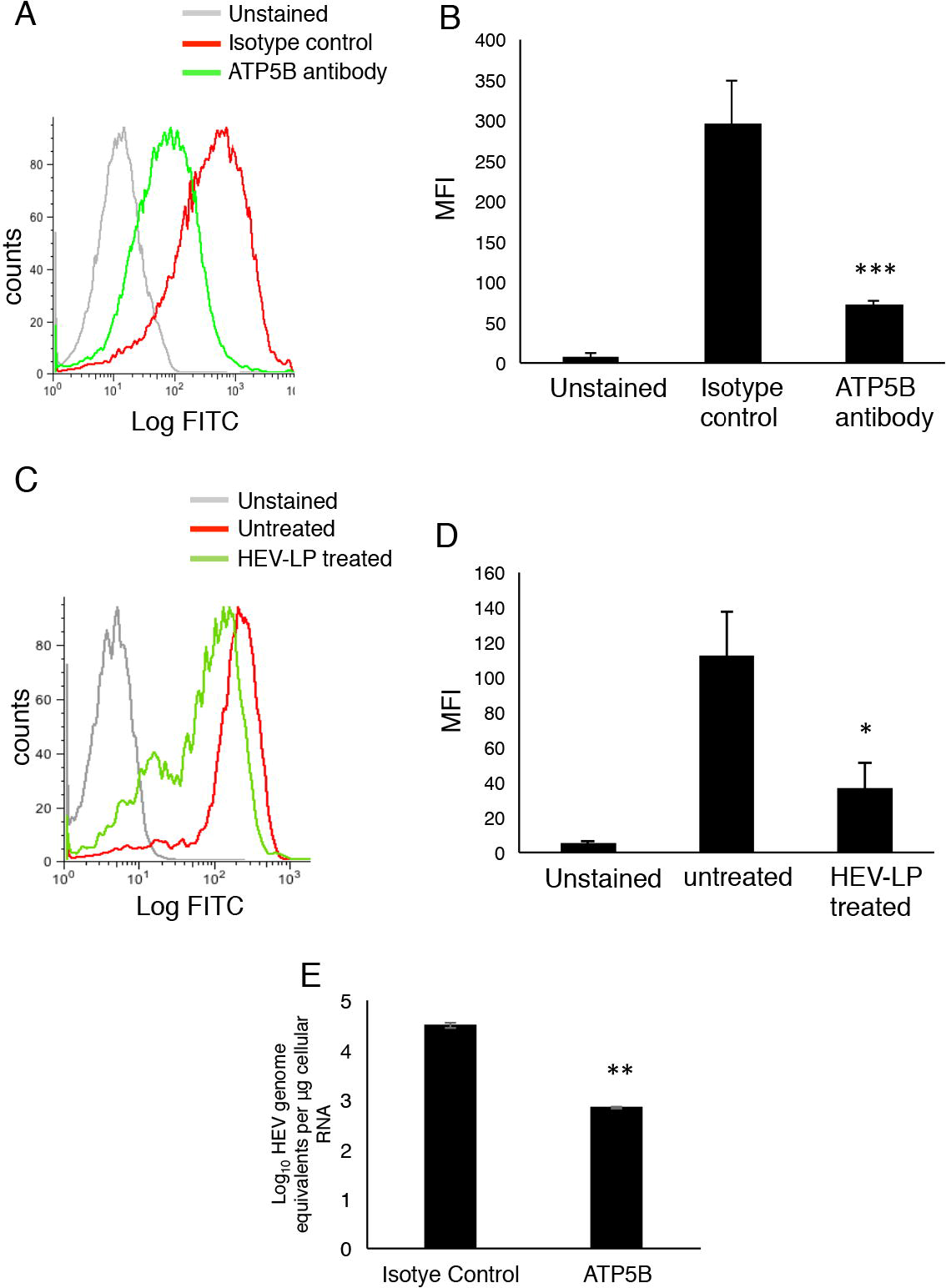
**(A)** Huh-7 cells were incubated with anti-ATP5B (or isotype control) antibodies, following which HEV-LP (50 μg) was bound to cells. Cells were stained with anti-HEV capsid antibody and subjected to flow cytometric analysis. Histograms were plotted using FlowJo software. **(B)** The mean fluorescence intensity from the previous experiment was plotted. Data represents average of three independent experiments ± SD ***p<0.005. **(C)** HEV-LP (50 μg) was bound to cells first, followed by staining for ATP5B. Flow cytometry was carried out and histograms were plotted using the FlowJo software. **(D)** Mean fluorescence intensity was plotted and data represents an average of three independent experiments ± SD *p<0.05. **(E)** HEV infection was carried out as described in Materials and Methods. Cells were incubated with anti-ATP5B (or isotype control) antibodies, HEV infection was carried out and 40 hr later HEV genome equivalents were estimated by qRT-PCR using a standard curve. Data are shown for three experiments ± SD (**p<0.01).

The binding of HEV-LP was also checked on cells transfected with siRNA against ATP5B. Knockdown of total cellular levels of ATP5B were checked in siRNA transfected cells by western blotting. There was about 5-fold and 1.5-fold reduction in ATP5B levels in Huh-7 and HEK293 cells, respectively (Fig. 3a). This was also reflected in the levels of ectopic ATP5B on surface staining of siRNA transfected cells with anti-ATP5B antibodies. In Huh-7 cells there was a significant reduction in ectopic ATP5B levels (Fig. 3b; p<0.01), and even though HEK293 cells also showed reduced ectopic ATP5B levels, the difference was not significant (Fig. 3c). As expected, HEV-LP binding was significantly reduced in Huh-7 and HEK293 cells that received ATP5B siRNA compared to scrambled siRNA transfected cells (Fig. 3d, 3e). The extent of inhibition was greater for Huh-7 cells compared to HEK293 cells. As in the case of antibody-mediated block, we wanted to test siRNA transfected cells for HEV infection. However, repeated attempts at this were unsuccessful. Unlike HEV-LP binding, the HEV infection protocol required siRNA transfected cells to be tested for virus two days after infection. This extended incubation of cells knocked down for an essential protein such as ATP synthase, made these cells non-viable.

**Figure 3.**
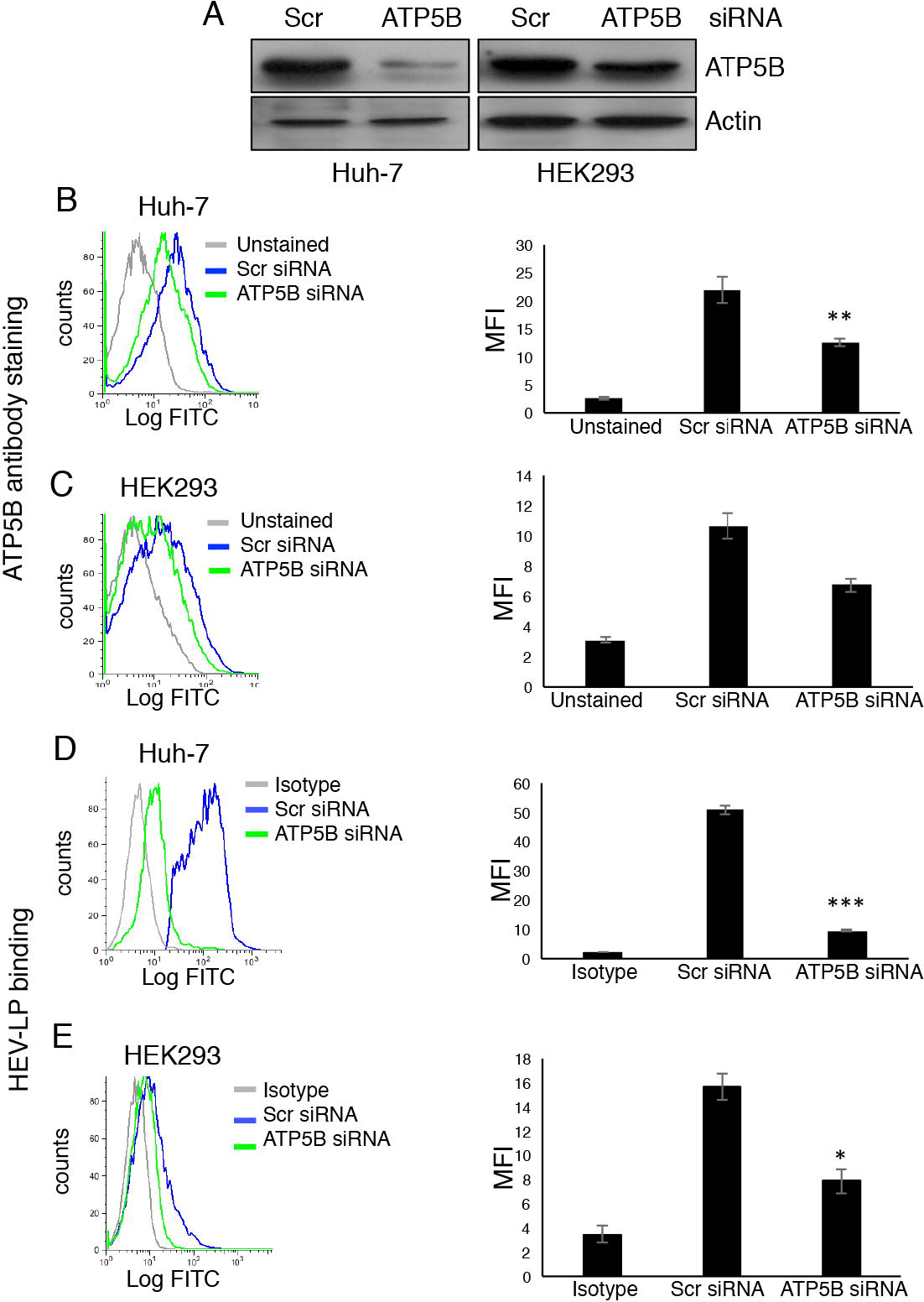
**(A)** Huh-7 or HEK293 cells were transfected with ATP5B-specific or scrambled control siRNAs as described in Materials and Methods. The extent of knockdown of total ATP5B in both cell lines was assayed by western blotting. Actin was used as a loading control. To check the knockdown of surface expressed ATP5B, Huh-7 cells **(B)** or HEK293 cells **(C)** were transfected with ATP5B-specific or scrambled control siRNA, fixed and stained with anti-ATP5B antibodies. Representative histograms plotted using FlowJo are shown on the left. Mean fluorescence intensity is shown on the right for both cell lines; data represents three independent experiments ± SD (**p<0.01). HEV-LP binding was checked in Huh-7 **(D)** or HEK293 **(E)** cells transfected with ATP5B-specific or scrambled control siRNAs. Representative histograms plotted using FlowJo are shown on the left and mean fluorescence intensities of three independent experiments ± SD are shown on the right for both cell lines (*p<0.05; ***p<0.001).

**Colocalization and intracellular trafficking of HEV-LP and ATP5B.** Viruses bind to surface receptors and are internalized along with the receptor in the form of a complex. We therefore asked whether cell surface bound HEV-LPs colocalized with ectopic ATP5B. For this, the HEV-LPs were labelled with FITC as described in Materials and Methods; these particles are called FITC-VLPs. Saturating concentrations of FITC-VLPs were bound to Huh-7 cells on ice for one hour (to prevent internalization of the FITC-VLP) and the cells fixed and counter-stained with anti-ATP5B antibody. The colocalization of surface bound FITC-VLPs was observed with cell surface ATP5B (Fig. 4a; n=22, Pearson’s coefficient=0.597). Alternatively, FITC-VLP bound cells were shifted to 37°C to allow internalization, and colocalization was checked at different times post-warming to determine if the internalized FITC-VLPs remained associated with ATP5B. For this, the cells were harvested at various times after FITC-VLP binding and internalization, surface stripped, fixed, permeabilized and stained with anti-ATP5B antibody. The internalized FITC-VLPs remained associated with ATP5B at 30 and 60 min, but the association was lost by 2 hours post-internalization (Fig. 4b, 4c, 4d). Binding of virus or HEV-LP to receptor molecules is likely to trigger internalization of the virus-receptor complex into internal compartments such as endosomes, and the internalized FITC-VLPs would be expected to remain associated with ATP5B up to a certain time, before dissociating from it.

**Figure 4.**
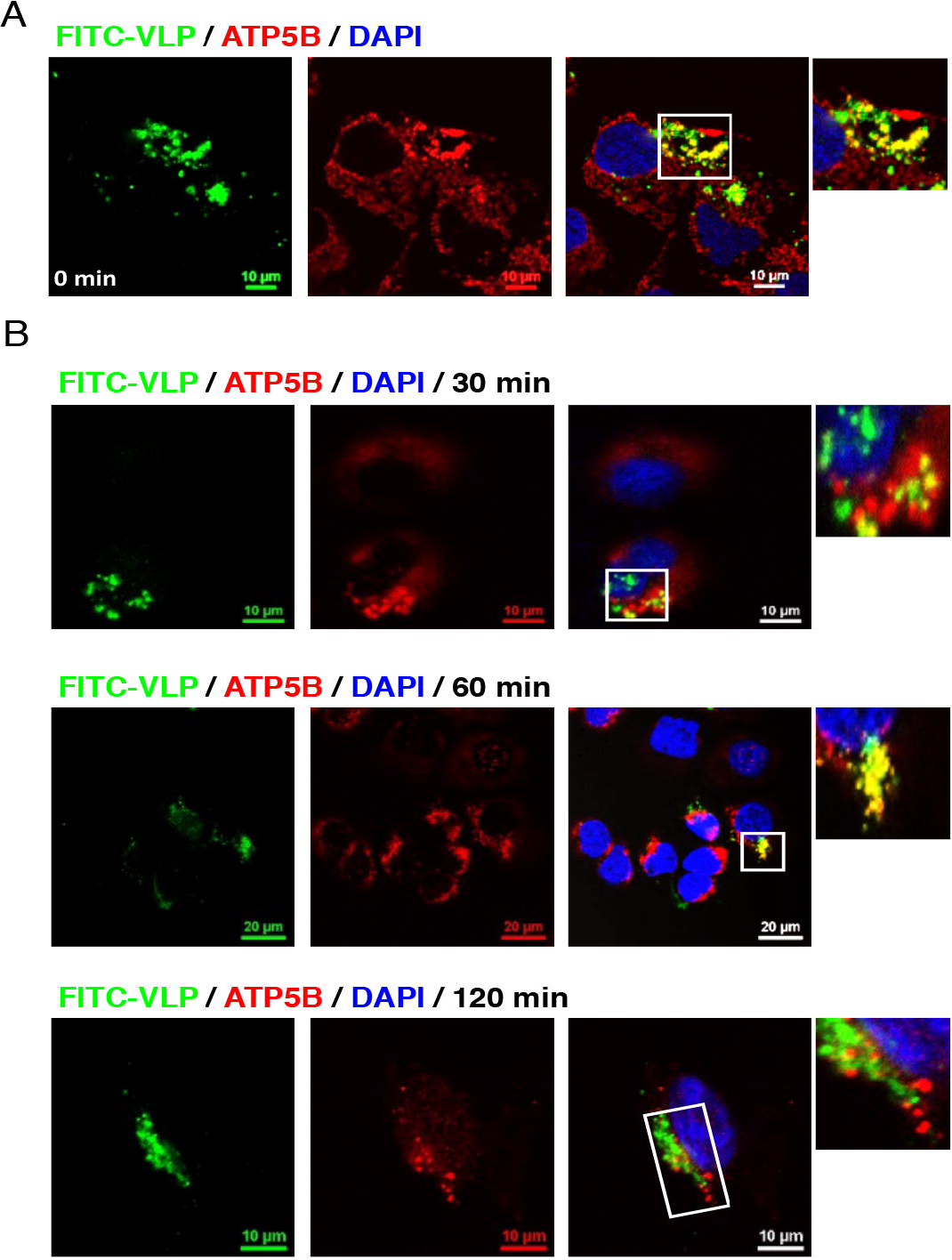
(Colocalization and trafficking of HEV-LPs and ATP5B.**(A)** FITC-VLP (green) was bound to Huh-7 cells on ice, the cells were fixed and counter-stained with anti-ATP5B antibodies (red) and visualized by confocal microscopy. **(B)** FITC-VLP (green) was bound to cells on ice and was subsequently allowed to internalize at 37°C. At the indicated times post-internalization, cells were fixed, permeabilized and stained for ATP5B (red), and visualized by confocal microscopy. The boxed regions have been enlarged on the right. Colocalization was quantified using the NIS Elements software. Scale bars are shown.

**Surface bound and internalized HEV-LPs are present in cholesterol-rich microdomains.** Ectopic ATP synthase has been shown to localize to cholesterol rich microdomains of the plasma membrane, also known as lipid rafts (Kim et al., 2004; Wang et al., 2006). These regions are enriched in molecules such as the GM1 ganglioside receptor and some integrins. Cholera toxin subunit B (CT×B) binds GM1 and is used as a marker for these membrane microdomains. To check if the HEV-LP was associated with lipid rafts during entry, we carried out binding of FITC-VLP and biotinylated CTxB to cells on ice for 1 hr, washed the cells and isolated lipid rafts using cold Triton extraction and sucrose density gradient centrifugation (Legler et al., 2005). Raft regions were detected by probing the fractions with Streptavidin-HRP. The surface bound FITC-VLP was found to partition to GM1-positive (raft enriched) as well as to EGFR-positive (non-raft enriched) fractions, although most of it was in the raft fractions (Fig. 5a). When FITC-VLP was allowed to internalize at 37°C for 1 hr after binding on ice, it remained associated with the GM1-positive fractions, although a significant portion was redistributed to the EGFR-positive non-raft regions (Fig. 5b). Since the VLP co-trafficks with ATP5B up to 1 hour post internalization (Figure 4), we conclude that some part of the internalized VLP remains associated with internalized ATP5B in the raft regions.

**Figure 5.**
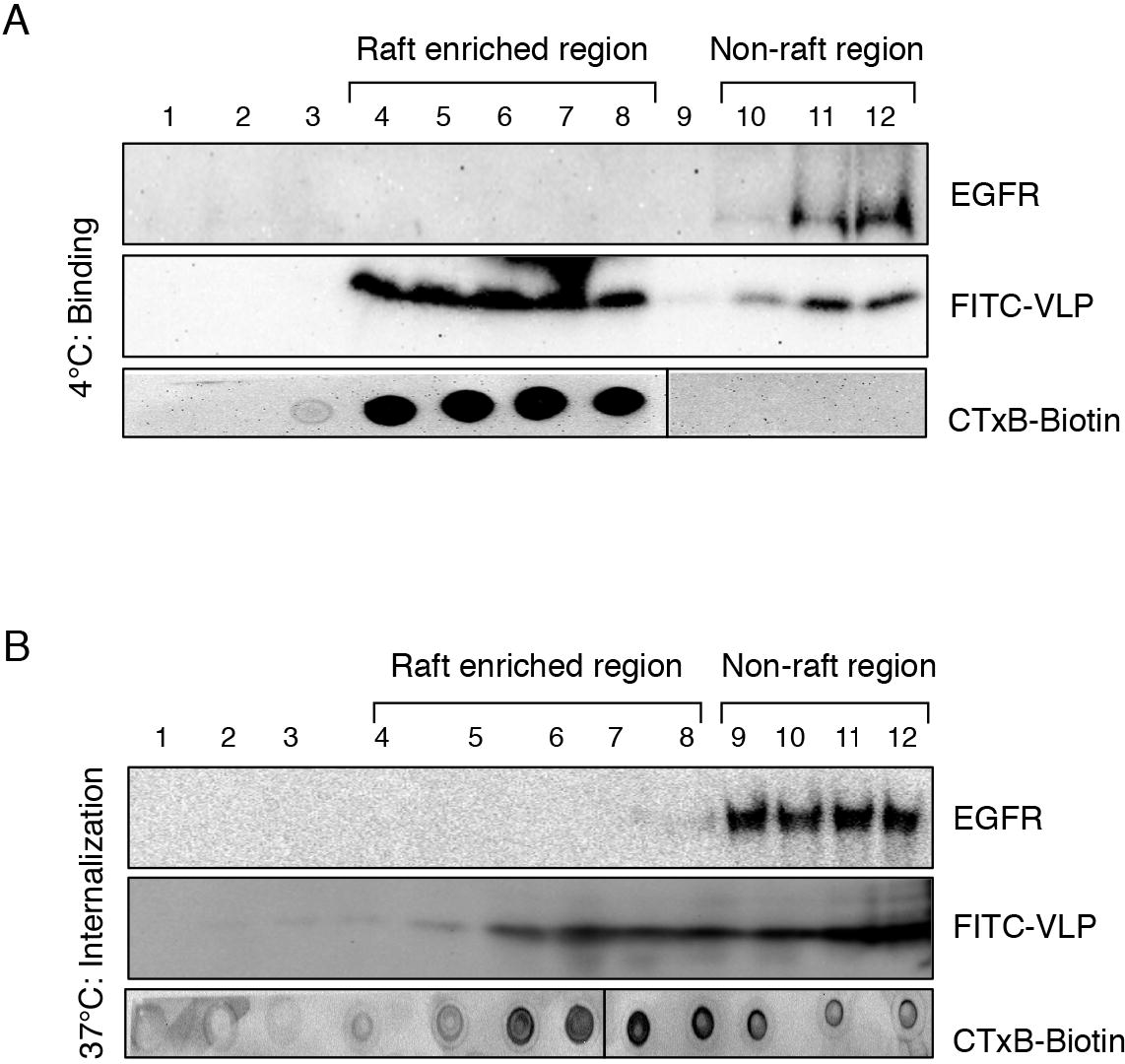
HEV-LPs bind lipid raft regions of the cell membrane. **(A)** FITC-VLP and biotinylated Cholera toxin subunit B (CTxB-Biotin) were allowed to bind to Huh-7 cells on ice for 1 hr, and cold Triton extraction was carried out followed by sucrose density gradient centrifugation as described in Materials and Methods. One mL fractions were collected from the top of the ultracentrifuge tube; 5 μL of each was blotted onto nitrocellulose membranes, blocked with BSA and probed with HRP-conjugated Streptavidin (bottom panel). Further, 50 μL from each fraction was run on a reducing SDS-polyacrylamide gel, transferred to nitrocellulose membrane and probed with antibodies to EGFR (a marker for non-raft membrane regions) or HEV-LP (ORF2 protein) as indicated. **(B)** FITC-VLP was allowed to internalize at 37°C after binding on ice for 1 hr, and lipid rafts were isolated and probed as described above.

This is further corroborated by recent studies from our laboratory (Holla et al., 2015), which show that reducing membrane cholesterol levels with Nystatin/Progesterone or Filipin also inhibited the entry of HEV-LPs and HEV infection of Huh-7 cells.

## DISCUSSION

Viruses bind to a diverse array of molecules on the surface of host cells to initiate infection. While some of these molecules are more generalized attachment factors that concentrate the virus on the cell surface, productive entry usually requires binding to one or more specific receptors. We have previously shown that HSPGs, specifically Syndecans, are critical attachment factors on liver cells that bind HEV (Kalia et al., 2009). However, the cellular receptor(s) for HEV, a virus that is endemic to large parts of the world and causes significant morbidity and mortality (Khuroo, 2011), have so far not been identified. The reasons for this and other lacunae in the understanding of HEV biology is its inability to grow efficiently in culture or to suitably infect small animals (Tanaka et al., 2007). We have circumvented this by utilizing a recombinant hepatitis E virus-like particle (HEV-LP) to identify potential cellular receptor(s), followed by confirmation with infectious virions.

Two cellular proteins, ATP synthase subunit β (ATP5B) and Ribophorin I (RPN1) were identified using a HEV-LP mediated pull down of plasma membrane proteins. The results were counter-intuitive since both of these molecules are not classically present on the cell surface. The ATP5B is part of the mitochondrial F_1_–F_0_ ATPase but is also found on the plasma membrane (Chi & Pizzo, 2006); RPN1 is part of the oligosachcharyltransferase complex in the rough endoplasmic reticulum (RER) and is involved in the N-glycosylation of proteins (Helenius & Aebi, 2004). Although RPN1 has not been reported to be present on the cell surface, our flow cytometric analyses and western blotting of plasma membrane fractions do indicate this (data not shown).

The ATP5B contains the catalytic core of the large multi-subunit ATP synthase complex, which drives ATP synthesis in the mitochondria (Boyer, 1997). Mitochondrial ATP synthase is translocated to cholesterol-rich ‘lipid raft’ regions of the plasma membrane, but the mechanism of this translocation is not understood (Kim et al., 2004). Further, adding cholesterol to HUVEC cells increased translocation of the ATP synthase to the plasma membrane (Wang et al., 2006). Surface localized or ‘ectopic’ ATP synthase is involved in the maintenance of intracellular pH, cholesterol homeostasis and cellular response to angiogenic factors (Chi & Pizzo, 2006), influenza virus maturation and budding (Gorai et al., 2012), and the transfer of HIV-1 particles from antigen presenting cells (APCs) to CD4+ T cells at the virological synapse (Yavlovich et al., 2012). Recently, ATP5B has also been shown to be involved in the entry of the Chikungunya virus (CHIKV) into insect cells (Fongsaran et al., 2014).

Since ectopic ATP synthase plays important roles in the entry and morphogenesis of other viruses, we investigated its role as an entry factor for HEV. Through a series of experiments that involved characterization of plasma membrane fractions, surface staining for ATP5B by flow cytometry, and mass spectrometric profiling of the surface proteome from different cell lines, we have shown that ATP5B is indeed present on the plasma membrane and its levels are higher in cells that are permissive for HEV infection compared to those that are not. However, it was interesting to find ATP5B in the p239-mediated pull down from both permissive (Huh-7) and non-permissive (HEK293) cells, although its levels were consistently higher in the former. This could be due to pull down using purified membrane fractions, where the concentrations of ATP5B were much higher as compared to whole cells. Additionally, the stoichiometry of binding of these membrane proteins to an immobilized VLP *in vitro* would be different from that on the cell surface. The permissive (Huh-7, PLC/PRF/5, A549) and non-permissive (HEK293) nature of cell lines might also be due to factors other than the presence or absence of a cell surface molecule that the virus uses for entry. For example, coxsackieviruses group B (CVB), bind CHO cells due to the presence of CAR receptors, although replication does not take place in these cells (Kramer et al., 1997) Similarly, most tissue macrophages are considered permissive for HIV-1 infection, since these are derived from monocytes. However, intestinal macrophages do not support HIV-1 infection. Exposure of monocytes to intestinal extracellular matrix (stroma)-conditioned media (S-CM) and HIV-1 simultaneously, revealed that the CD4+ CCR5+ subset of differentiated intestinal macrophages did not support HIV-1 infection, indicating that factors other than receptor binding may be involved in making cells permissive or otherwise (Shen et al., 2011).

Huh-7 cells pre-incubated with antibodies against ATP5B showed reduced HEV-LP binding; conversely, surface staining for ATP5B was also significantly reduced when HEV-LP was pre-bound to cells. The knockdown of ATP5B with siRNA also led to reduced HEV-LP binding to Huh-7 cells. During internalization of surface-bound HEV-LPs, these were found to colocalize with ATP5B for up to one hour postinternalization, following which this association was lost. The internalized HEV-LPs also colocalized with CT×B, which binds the GM1 receptor and is a marker for cholesterol-enriched membrane microdomains or ‘lipid rafts’ (Torgersen et al., 2001). This was also the case when these membrane microdomains were prepared using cold Triton extraction and sucrose gradient centrifugation. This could be due to binding of the HEV-LPs to the ectopic ATP synthase complex, which resides in these membrane fractions (Kim et al., 2004). To validate our findings, surface exposed ATP5B was blocked with specific antibodies and the cells were infected with virions from the stool of an infected hepatitis E patient. Intracellular virus replication was compromised in Huh-7 cells pre-treated with anti-ATP5B antibodies. This reduced replication is likely to be due compromised HEV binding and internalization, and validates our findings that ATP5B is involved in the binding and entry of HEV into liver cells.

## ACKNOWLEDGEMENTS

This work was supported by a grant (to SJ) from the Department of Biotechnology, Government of India and Research Fellowships (to PH and IA) from the Council for Scientfic and Industrial Research, India.

